# The *Xanthomonas fragariae* effector XopK suppresses stomatal immunity by perturbing abscisic acid accumulation and signaling pathways in strawberry

**DOI:** 10.1101/2024.02.07.579334

**Authors:** Xiao-lin Cai, Wenyao Zhang, Haiyan Yu, Ying-qiang Wen, Jia-yue Feng

**Affiliations:** State Key Laboratory of Crop Stress Biology for Arid Areas, College of Horticulture, Northwest A&F University, Yangling, Shaanxi, China; Key Laboratory of Protected Horticulture Engineering in Northwest China, Ministry of Agriculture, Yangling, Shaanxi, China

**Keywords:** Strawberry, *Xanthomonas fragariae*, stomatal immunity, abscisic acid, methyl jasmonate

## Abstract

*Xanthomonas fragariae* (*Xaf*) usually causes angular leaf spot (ALS) in strawberry all over the world. Recently, we isolated a new strain of *Xaf* called *Xaf* YL19. This strain causes not only typical ALS symptoms, but also dry cavity rot in the crown tissues of strawberries. This was the first time a *Xaf* strain had both of these effects in strawberries. Pathogen effectors play a crucial role in plant colonization and infection. Understanding how effector proteins escape from plant surveillance is important for plant breeding and resistance deployment. In this study, thirty-three putative effectors were predicted in *Xaf* YL19, and a virulent secreted effector called XopK was identified. XopK has a robust capacity for suppressing plant cell death and transcript response to infection host. Transgenic strawberries expressing XopK exhibit more susceptibility to *Xaf* YL19, and this was associated with weakened stomatal immunity. Additionally abscisic acid (ABA) accumulation and signaling were mainly suppressed in the presence of XopK. XopK also inhibited ABA- and methyl jasmonate (MeJA)-induced stomatal closure, and ABA- but not MeJA-induced reactive oxygen species (ROS) production. Moreover, endogenous ABA is critical for *Xaf*-induced ROS burst and stomatal closure. These results suggested that *Xaf* YL19 uses XopK to suppress JA, especially ABA signaling, for disrupting stomatal closure, and allowing its colonization for disease development. Taken together, our results provide a novel virulent mechanism by *Xaf* and an available strategy for resistant breeding in strawberries.

## 1. Introduction

Strawberry (*Fragaria* × *ananassa*) is a very valuable fruit crop that is extensively grown worldwide. In 2020, its global output was estimated to be worth 14 billion USD (Hernández-Martínez et al., 2023). *Xanthomonas fragariae* (*Xaf*) causes bacterial angular leaf spot (ALS) in strawberries, which is one of the most severe diseases for this fruit crop (EPPO, 2023; Kim et al., 2016; Roach et al., 2016; Wang & Turechek, 2016). Recently, we isolated a *Xaf* new strain, YL19, which was observed to cause both typical ALS symptoms in leaves and dry cavity rot in the crown of strawberries, the first *Xaf* strain has both these effects in strawberries (Li et al., 2021; Wang et al., 2023). The pathogen usually enters through the leaf stomata and colonizes the parenchyma apoplast. Initially, it causes water-soaked spots on the leaves which later turn translucent and contiguous patches of lesions. The pathogen then spreads systematically to other parts of the plant, including the crowns and runners, even if no apparent symptoms are present (Luiz et al., 2022; EPPO, 2023; Bestfleisch et al., 2015; Wang et al., 2023). Thus, it is essential for studying how *Xaf* infect strawberries through stomata to develop effective management strategies.

When plants are infected by a pathogen, their guard cells respond quickly by lowering their turgor pressure, which leads to the closure of stomata, a process called stomatal immunity (Wu & Liu, 2022). The early stages of bacterial infection are greatly limited by stomatal immunity (Melotto et al., 2017). Stomatal closure is often triggered by increased various second messengers, such as reactive oxygen species (ROS), nitric oxide (NO), and cytosolic Ca^2+^ in guard cells. These second messengers are induced by phytohormone signaling, such as abscisic acid (ABA), jasmonates (JA) and salicylic acid (SA) (Munemasa et al., 2011). As a counteracting strategy, most phytopathogenic bacteria, including *Xanthomonas* spp., rely on type III effectors (T3Es) as important weapons to actively induce stomatal reopening for the benefit of infection. They do this by altering phytohormones levels or signaling pathways (Melotto et al., 2017). Like XopC2 from *Xanthomonas oryzae* pv. *oryzicola* (*Xoc*) RS105 (Wang et al., 2021), HopX1 from *Pseudomonas syringae* pv. *tabaci* (*Pta*) 11528 (Gimenez-Ibanez et al., 2014), both of them promote the degradation of JAZ proteins to activate JA signaling for inhibiting stomatal immunity in rice and *Arabidopsis*, respectively. Elevated ABA levels manipulated by effectors, such as AvrXccC8004 from *X. campestris* pv. *campestris* (Ho et al., 2013), AvrPto, and AvrPtoB from *P. syringae* DC3000 (de Torres-Zabala et al., 2007), have been associated with *Arabidopsis* infections. Still, the exact molecular mechanisms behind the majority of effector-induced stomatal reopening are yet unknown.

The genomic information of 74 *Xaf* strains, including the new strain YL19 (Li et al., 2021; Wang et al., 2023; Wei et al., 2022; Qiu et al., 2023), was sequentially made accessible to the public (Gétaz et al., 2017; Henry & Leveau, 2016; Vandroemme et al., 2013; Qiu et al., 2023). The virulent *Xaf* strain LMG25863 was predicted to contain twenty-three T3Es and several putative new effectors (Vandroemme et al., 2013). Furthermore, transcriptome analysis of *Xaf* in strawberries revealed that T3Es can be functional in *Xaf* during the pathogenesis process (Gétaz et al., 2020; Puławska et al., 2020). Recently, 47 T3Es were mined based on a machine-learning method, and two novel *Xaf* T3Es, XopBG and XopBF, were translocated and elicited hypersensitive reaction (HR) on pepper leaves, revealed its possible pathogenic role in *Xaf* Fap21 (Wagner et al., 2023). So far, there is still very limited knowledge on the in-depth interaction mechanism between *Xaf* T3Es and strawberries, which may be attributed to the difficulties in overcoming technological hurdles associated with *Xaf* genetic editing and the absence of resistant materials (Vandroemme et al., 2013; Maas et al., 2000, 2002; Roach et al., 2016). Thus, exploring the key virulent T3Es of *Xaf* and revealing the mechanism behind the interaction between *Xaf* and strawberry is essential for the prevention of disease.

In this study, we examined the capacity of triggering or inhibiting cell death, and transcript level during infection, in 22 of thirty-three putative effectors of *Xaf* YL19. We identified a virulent effector XopK, and ectopically overexpressed XopK in strawberries to assess its function. Further, we found that XopK disrupts ABA and JA signaling to favor pathogen infection. This study provides a novel virulent mechanism by *Xaf* and an available strategy for resistant breeding in strawberries.

## 2 Results

### 2.1 XopK is a potential virulent effector in *X. fragariae* YL19

To mine virulent T3Es in the strain YL19 of *X. fragariae*, we predicted thirty-three T3Es in the whole genome of *Xaf* YL19 based on the homolog blast (identity ≥40%, query length ≥75% in nucleotide sequence) in Xanthomonads species (Fig. S1a; Table S1). Three T3Es including XopK, XopQ, and XopZ2 are very conserved in all 21 test strains of the ten *Xanthomonas* genus, while other T3Es distribute variable (Fig. S1b). Then, we carried out a preliminary assessment for 22 of thirty-three T3E functions by investigating the capacity of inducing or suppressing cell death in *N. benthamiana*. Five effectors, namely, XopD, XopAE, XopF1, XopF2, and XopX2, were observed that could trigger HR at five days after infiltration (dai). Then, the remaining 17 non-HR effectors were expressed in *N.benthamiana*, and the BAX (a cell death elicitor) infiltrated later at 24-hour intervals as described in Figure S3-a. At 5 dai, five effectors including XopK, XopQ, AvrBs1, XopE4, and XopAF completely suppressed BAX-induced cell death, and other effectors with weakened or without suppression activity (Fig. S3b). Furthermore, considering the beginning of ALS and the bacterial population significantly increased (*P*=0.0361) at 96 h post-inoculation (hpi) (Fig. S4a, b), the transcript levels of those twenty-two T3Es in *Xaf* YL19-infected strawberry plants were determined over a 96-h course. Compared to 0 hpi, thirteen T3Es kept low expression. They were even suppressed at 54 hpi, and only nine effectors (XopH, XopC1, XopZ2, XopAF, XopE5, XopC2, XopK, XopB, and XopV) were highly expressed at 54 or 96 hpi (Fig. S4c), indicating those effectors may play important role in the early infection. Taken together, the effector XopK was selected as a key potential virulent effector for further study because of its conservation, robust capacity to suppress cell death, as well as transcripts response to colonize host cells.

### 2.2 XopK could be translocated into the host cell and localize plasma-membrane

To detect whether XopK protein could be translocated into host cells during infection, a phiLOV2.1 fluorescence protein was fused with XopK under the control of its native promoter (Roushan et al., 2018), then cloned into the pBBR5 vector and transformed into *Xaf* YL19 competent cells, hereafter named *Xaf* YL19[*XopK:phiLOV2.1*]. The strain, *Xaf* YL19[*phiLOV2.1*], expressing *phiLOV2.1* under the same promoter of XopK as control (Fig. 1a). The leaves and roots of strawberry were inoculated with *Xaf* YL19[*XopK:phiLOV2.1*] and *Xaf* YL19[*phiLOV2.1*] for five days, then the phiLOV2.1 fluorescence signal were observed using the confocal microscopy. A bright filamentous structure or a dot-shaped green fluorescence was observed in leaves or roots, respectively, in the inoculation of *Xaf* YL19[*XopK:phiLOV2.1*] but not that of *Xaf* YL19[*phiLOV2.1*]*-*inoculated plants (Fig. 1b). The results indicated that XopK could be translocated into the host cell by *Xaf* YL19.

**Figure 1.**
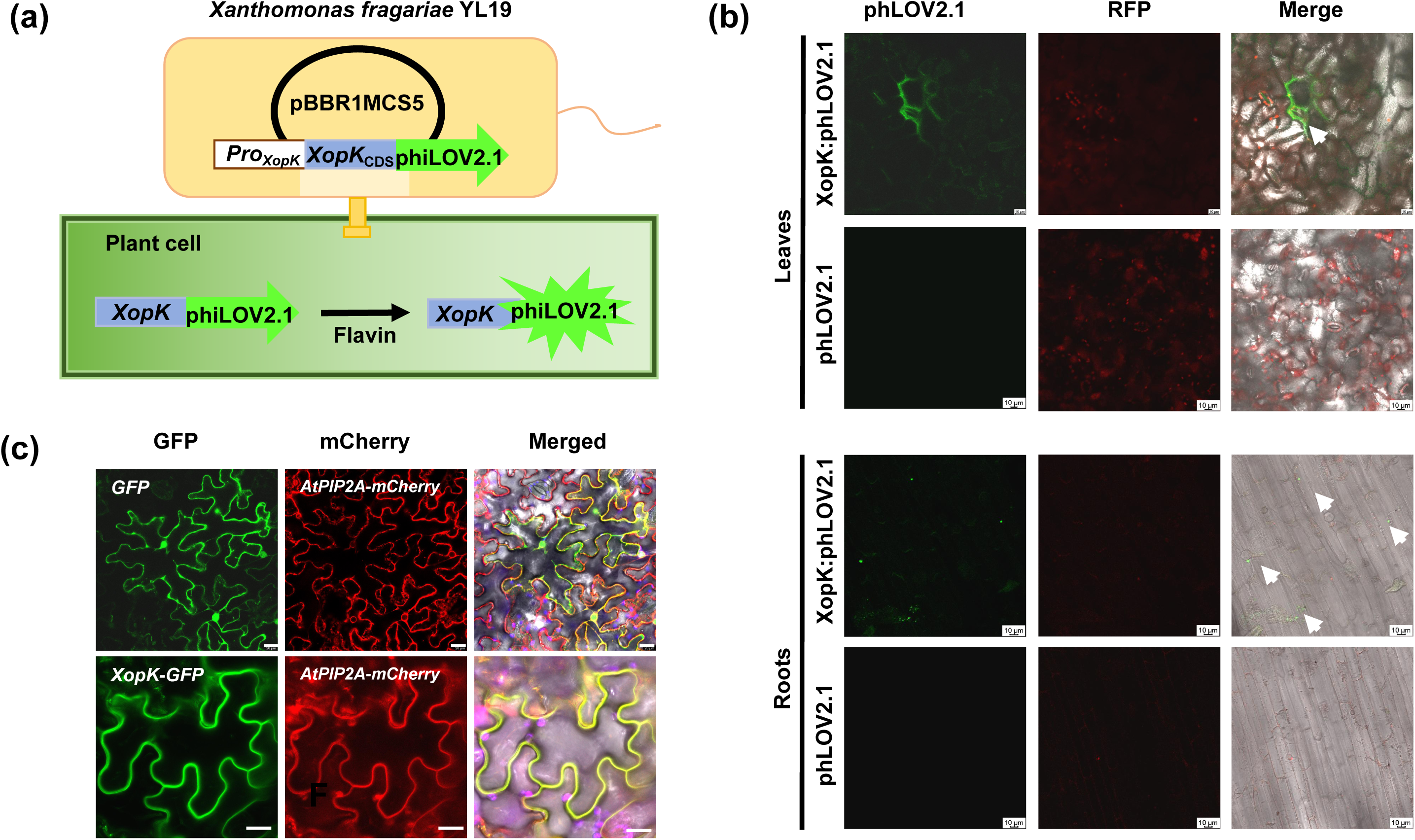
XopK is a secreted effector and localizes in plasma membrane. (a) Translocation assay schematic diagram. XopK fused with fluorescence protein phiLOV2.1 under the control of XopK native promoter (960 bp) was cloned into pBBR5 vector, then the recombination plasmid transformed into the competence of Xaf YL19. (b) Translocation of phiLOV2.1-tagged XopK proteins from Xaf YL19[XopK:phiLOV2.1] to strawberry roots and leaves. Images were captured by confocal microscopy three days after inoculation. The fluorescence of phiLOV2.1 was excited at 488 nm. Scale bars: 20μm. (c) Subcellular localization of XopK in N.benthomiana. mCherry-tagged AtPIP2A as a membrane-localized marker. Fluorescence of GFP and mCherry was excited at 488 nm and 561 nm, respectively.

XopK does not contain any known structural or functional domain predicted from its amino acid sequence via SMART and Pfam searches (Letunic et al., 2021; Mistry et al., 2021). However, it featured four transmembrane regions (predicted by TMHMM-2.0) (Fig. S5), which suggests that it is a protein associated with membranes. To experimentally confirm the location, *N. benthamiana* co-expressed the GFP-fusion XopK protein with mCherry-tagged AtPIP2A protein that is known situated on the membrane (Pumplin & Harrison, 2009). As seen in Figure 1(c), the GFP was diffuse throughout the plant cytoplasm and nucleus. XopK-GFP appeared to be associated with the plasma membrane (PM), which a visible overlap fluoresence signal with AtPIP2A-mCherry, suggesting that XopK localizes the plasma membrane in *N. benthamiana*.

### 2.3 Overexpression of XopK causes strawberry-enhanced susceptibility to *X. fragariae* YL19

To investigate how XopK functions in strawberry plants, we generated two independent transgenic strawberry lines expressing *XopK* in the background of *Fragaria vesca* ‘Hawaii 4’ (named *XopK*-OX1 and *XopK*-OX2, respectively). The protein abundance of GFP-fused XopK in *XopK*-OX lines was confirmed (Fig. S6a). No apparent differences in growth phenotype were observed between the wild-type and *XopK*-OX strawberries under normal growth conditions (Fig. S6b). Then, to assess the resistant level of *XopK* transgenic strawberries, *Xaf* YL19 suspension (10^8^ cfu/mL) was sprayed on strawberry leaves. Compared to the wild type, both two *XopK*-OX lines showed much more angular spots in leaves at 4 dpi and 8 dpi (Fig. 2a). The mean leaf disease index reached 62.7% and 60.5% in *XopK*-OX1 and *XopK*-OX2 plants, significantly higher (*P*=0.0070, and 0.0192, respectively) than that of the wild type (51.6%) (Fig. 2b). Consistently, in strawberry crowns, the wild type exhibited only mild water-soaked symptoms and slight brown spots, but more severe water-soaked symptoms and some even had small cavities were observed in *XopK*-OX lines at 45 dpi (Fig. 2a). The incidence of dry cavity rot in crown of *XopK*-OX1 (*P*=0.0078) and *XopK*-OX1 (*P*=0.0314) were also significantly higher than that of the wild type at 45 dpi (Fig.2c). In addition, bacterial population in the *XopK*-OX strawberry leaves were significantly (*P*<0.05) higher than the wild type at 4 dpi but not at 8 dpi (Fig. 2d), suggesting a possibility of XopK contributes to the initial infection of *Xaf* YL19. The hypothesis was also supported by the fact that there were no significant differences in both disease symptoms and bacterial population by two other inoculation ways including pressure-inoculation in leaves or syringe-inoculation in crowns (Figs. S7b, c).

**Figure 2.**
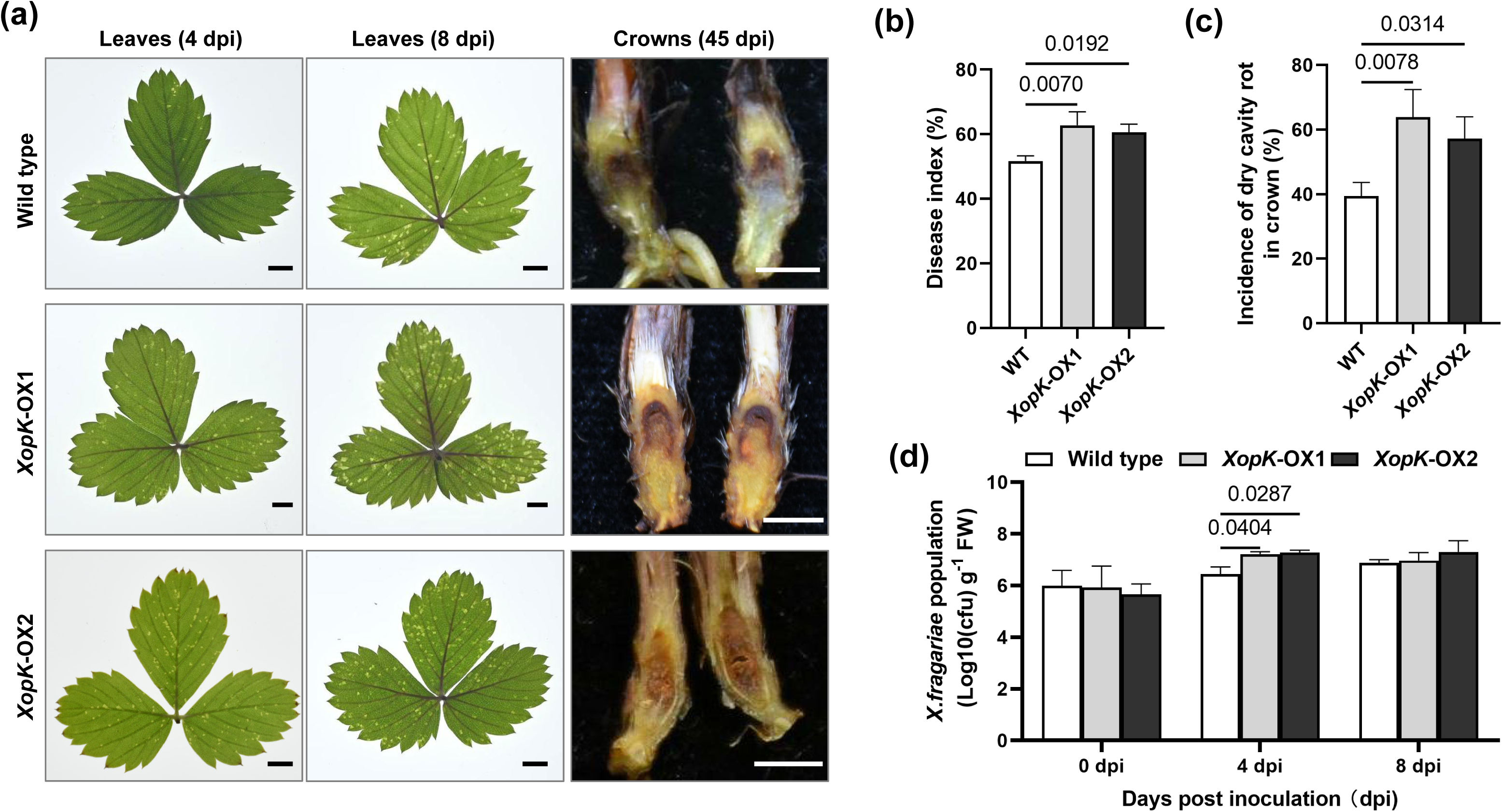
Overexpression of XopK causes strawberry-enhanced susceptibility to X.fragariae YL19. (a) Disease symptoms were photographed at 4 and 8 days post-inoculation (dpi) for leaves and at 45 dpi for crowns, respectively; Bars scale, 50 mm. Disease index (b), and incidence of dry cavity rot in the crown (c) of the wild type and XopK-OX strawberry lines. Data are mean ± SD (n=3). (d) Xaf YL19 population in leaves of two independent XopK-OX strawberry lines was determined at 0, 4, and 8 dpi by RT-qPCR. CFU, colony-forming unit. Data are mean ± SD (n=4). The values on the top of each column indicate a significant difference by Two-way ANOVA (Dunnett’s multiple comparisons test).

### 2.4 XopK suppresses pathogen-triggered stomatal closure and compromises drought tolerance in strawberry

Considering stomata are the main invading entry by *Xaf* on unwounded strawberry leaves, stomatal conductance (Gs) was evaluated after surface inoculation with *Xaf* YL19 bacterial suspensions. As shown in Figure 3(a), similar Gs were exhibited in the mock (10 mM MgCl_2_) treatment group, while Gs were significantly higher in the *XopK*-OX strawberry lines compared to the wild type at 2 h post-inoculation, suggesting XopK suppresses stomatal closure when *Xaf* YL19 is invading. The stomatal aperture with the same treatment also supports the conclusion (Fig. 3b). However, this difference is unrelated to average stomatal density (Fig. 3c). In addition, the water loss for detached leaves of *XopK*-OX strawberry lines was more rapid than that of wild type over the 40 min course (Fig. 3d). These results indicate that XopK suppresses *Xaf*-triggered stomatal closure and compromises drought-stress tolerance in strawberry.

**Figure 3.**
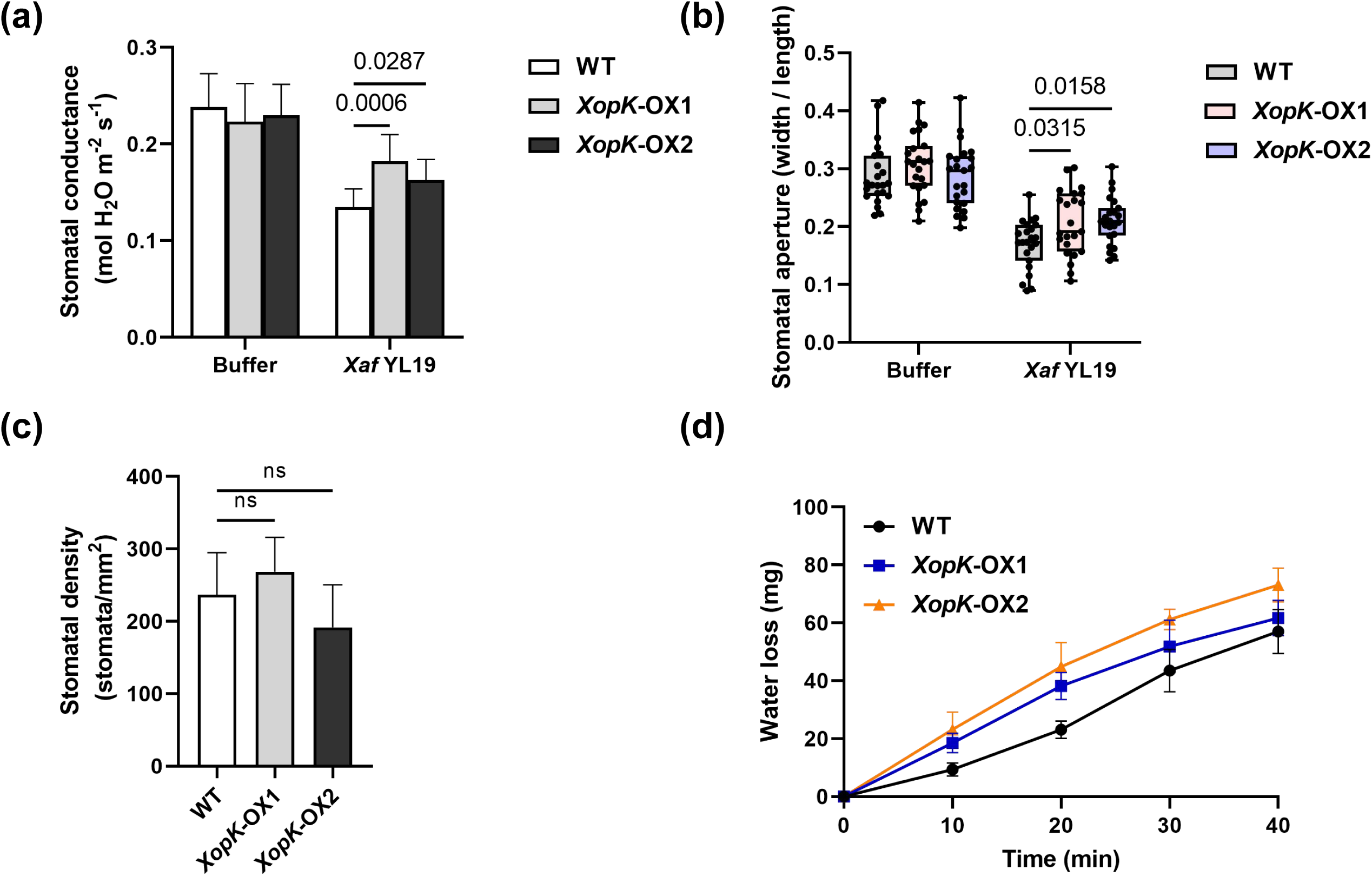
XopK causes defective of X.fragariae-induced stomatal closure and insensitive to drought stress in strawberry. Stomatal conductance (Gs) (a), data are mean ± SD (n=9), and stomatal aperture (b) of wild type and XopK-OX strawberry lines at 2 hours post-inoculation (hpi) with X.fragariae YL19 suspension. They were spraying equal volume buffer (10 mM MgCl_2_) as control. 23 stomata were measurement in each treatment. (c) Stomatal density of wild type and XopK-OX strawberry lines. Data are mean ± SD (n=8). (g) Water loss of detached leaves of wild type and XopK-OX strawberry lines ever<u>y</u> 10 min in a growth chamber. Data are mean ± SD (n=5). The values on the top of each column indicate a significant difference by Two-way ANOVA (Dunnett’s multiple comparisons test), ns, no significant (P>0.05).

### 2.5 Impairment of endogenous ABA accumulation and signaling in the presence of XopK in strawberry

The above results let us speculate that the phytohormone ABA, which often mediates drought stress responses and parallelly defends against pathogens by regulating stomatal closure, may be impaired in XopK-expressing strawberries. We thus examined the accumulation of ABA in *XopK*-OX strawberry lines. The results showed that endogenous ABA levels in *XopK*-OX1 and -OX2 strawberry leaves were about 73% and 47% of those wild-type plants (Fig. 4a). Consistently, the *9-cis-epoxy carotenoid dioxygenase 5* (*NCED5*), which encodes the rate-limiting enzyme for ABA biosynthesis (Zhou et al., 2021), was more transcriptionally suppressed in *XopK*-OX strawberry (Fig. 4b). In addition, the transcript level of *abscisic acid-insensitive 5* (*ABI5*) and *sucrose nonfermenting 1-relative protein kinase 2.6* (*SnRK2.6*) was significantly (*P*<0.05) reduced (Figs. 4c, d). In contrast, *XopK*-OX lines exhibited a similar level of both the JA and SA with the wild type (Fig.S7b, S8c). In addition, the expression of two JA-response genes including *JASMONATE ZIM-domain 1* (*FveJAZ1*) and *Lipoxygenase 14* (*FveLOX14*) was mild change or suppressed in the *XopK*-OX strawberry, respectively (Figs. S7c, d). These results revealed that XopK ectopically expression specifically suppresses ABA biosynthesis and signaling in strawberries.

**Figure 4.**
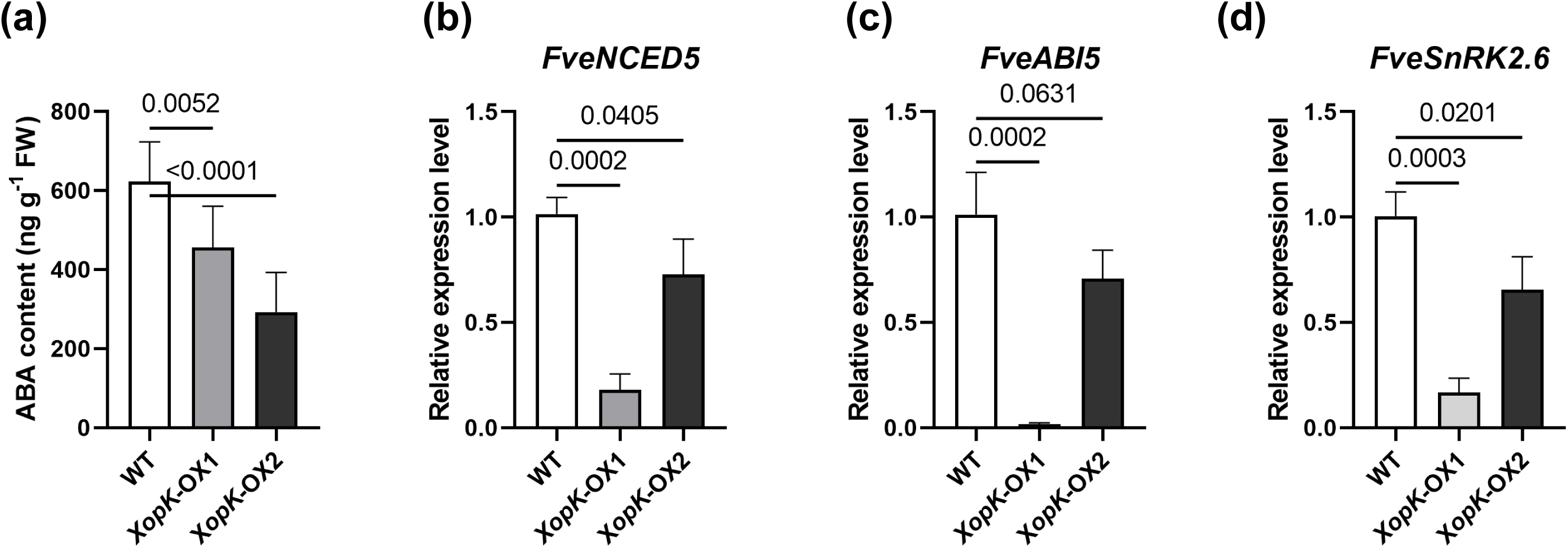
Impairment of abscisic acid (ABA) accumulation and signaling in the presence of XopK in strawberry. (a) Endogenous ABA level in wild type and *XopK*-OX strawberry under normal growth conditions. Data are mean ± SD (n=8). The expression level of *FveNCED5* (b), *FveABI5* (c) and *FveSnRK2.6* (d) in wild type and *XopK*-OX strawberry under normal growth conditions. Data are mean ± SD (n=3). *FveGAPDH* was used as the internal control. The values on the top of each column indicate a significant difference by Two-way ANOVA (Dunnett’s multiple comparisons test).

### 2.6 XopK suppresses ABA- and MeJA-induced stomatal closure

To clarify the defected-ABA level and signaling whether affects ABA- and MeJA-induced stomatal closing in *XopK*-OX strawberries. The time course of stomatal closing by application of 10 μM ABA was shown in Figure 5(a, b), the wild type reduced significantly (*P*=0.0004) stomatal apertures by 29% at 20 min, while the stomatal aperture of *XopK*-OX1 and *XopK*-OX2 exhibited only 0.09% and 0.12% closing at 60 min, respectively, but clear closing at 90 min. Strikingly, we also found 75 μM MeJA, even concentration of up to 200 μM (Fig.S7a), failed to induce stomatal closure in both two *XopK*-OX lines throughout the two-hour time duration, while stomata closed by 31% at 10 min (P<0.0001) and almost closing at 40 min after MeJA treatment for the wild type (Figs. 5c, d). In addition, the application of 30 μM SA induced stomatal closure at 10 min (Figs. S8a, b). Collectively, these results showed that ABA-, especially MeJA-, but not SA-induced stomatal closure is mainly defective in the presence of XopK in strawberries.

**Figure 5.**
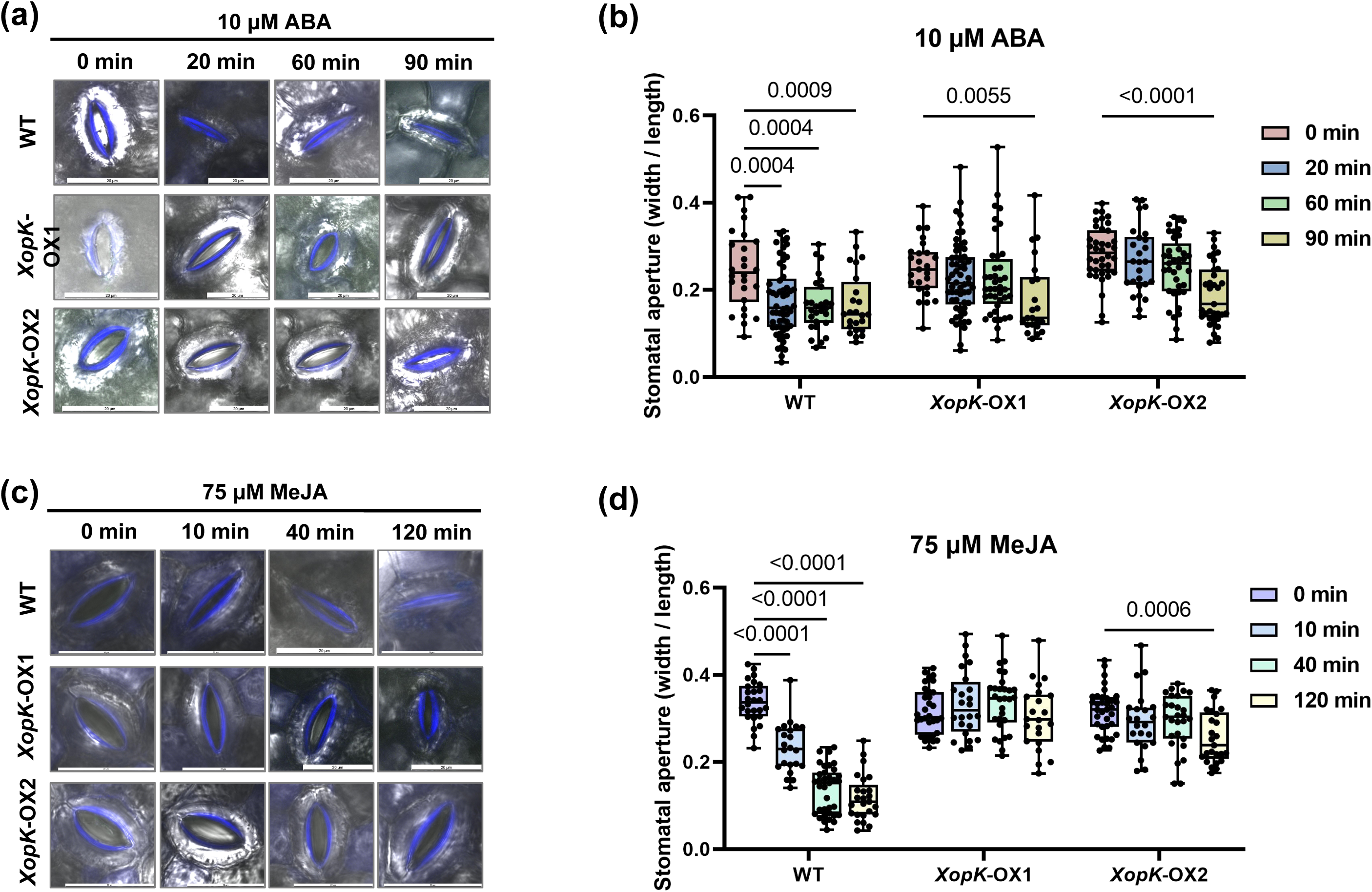
XopK suppresses ABA- and methyl jasmonates (MeJA)-induced stomatal closure in strawberries. Representative images of stomatal aperture (a), and the size of stomatal aperture (b) with the treatment of 10 μM ABA in wild type and *XopK*-OX strawberry over 90 minutes. Representative images of stomatal aperture (c), and the size of stomatal aperture (d) with the treatment of 75 μM MeJA in wild type and *XopK*-OX strawberry over 120 minutes. At least 25 stomatal apertures were counted at each time point. Scale bar 20 μm. The values on the top of each column indicate a significant difference by Two-way ANOVA (Dunnett’s multiple comparisons test).

### 2.7 Endogenous ABA is critical for *Xaf*-induced ROS burst and stomatal closure in strawberry

ROS accumulation is often observed downstream of ABA and MeJA in the signaling cascade leading to stomatal closure (Song et al., 2014). To further clarify the signaling events modulated by XopK, ABA- and MeJA-induced ROS levels in *XopK*-OX lines were analyzed with diaminobenzidine tetrahydrochloride (DAB) staining and nitroblue tetrazolium (NBT) staining. Application of 10 μM ABA and 75 μM MeJA significantly (*P*<0.001) induced ROS production in the wild type, whereas a significant defective accumulation of ROS after ABA but not MeJA treatment in leaves of *XopK*-OX lines (Figs. S9), suggesting that XopK may mainly suppress ABA- but not MeJA-induced ROS production.

In addition, we further used exogenous ABA and fluridone (FLU, an ABA biosynthesis inhibitor) to verify the function of ABA in *Xaf*-induced ROS bursts and stomatal closure. Pretreating with FLU at 100 μM caused a significantly higher staining intensity of DAB (*P*<0.0001) and NTB (*P*=0.0005) in the wild-type leaves and larger (*P*=0.0240) stomatal aperture than the control (10 mM MgCl_2_), whereas FLU alone did not make a significant difference (*P*>0.05) in both DAB and NBT staining intensity and stomatal aperture (Figs. 6a, b, c). In addition, the application of 10 μM ABA for 3 h in *XopK*-OX leaves followed by *Xaf* YL19 spraying-inoculation resulted in an increase in ROS production (1.57-fold in DAB staining intensity and 2.78-fold in NBT staining intensity) and smaller stomatal aperture in comparison to the control treatment without ABA (Figs. 6e, f, g). Consistently, FLU-pretreating significantly (*P*=0.0311) increased the bacterial population than the mock, and ABA-pretreating compensated *XopK*-expressing strawberry resistance to *Xaf* YL19. Notably, FLU did suppress ABA accumulation in wild type and exogenous ABA also did promote expression of *FveABI5 and FveSnRK2.6* in *XopK*-OX strawberry (Figs, S11, a, b, c), but 10 μM ABA and 100 μM fluridone did not inhibit the growth of *Xaf* YL19 (Fig. S11d). These results indicated that endogenous ABA is largely responsible for initial infection by regulating ROS production.

**Figure 6.**
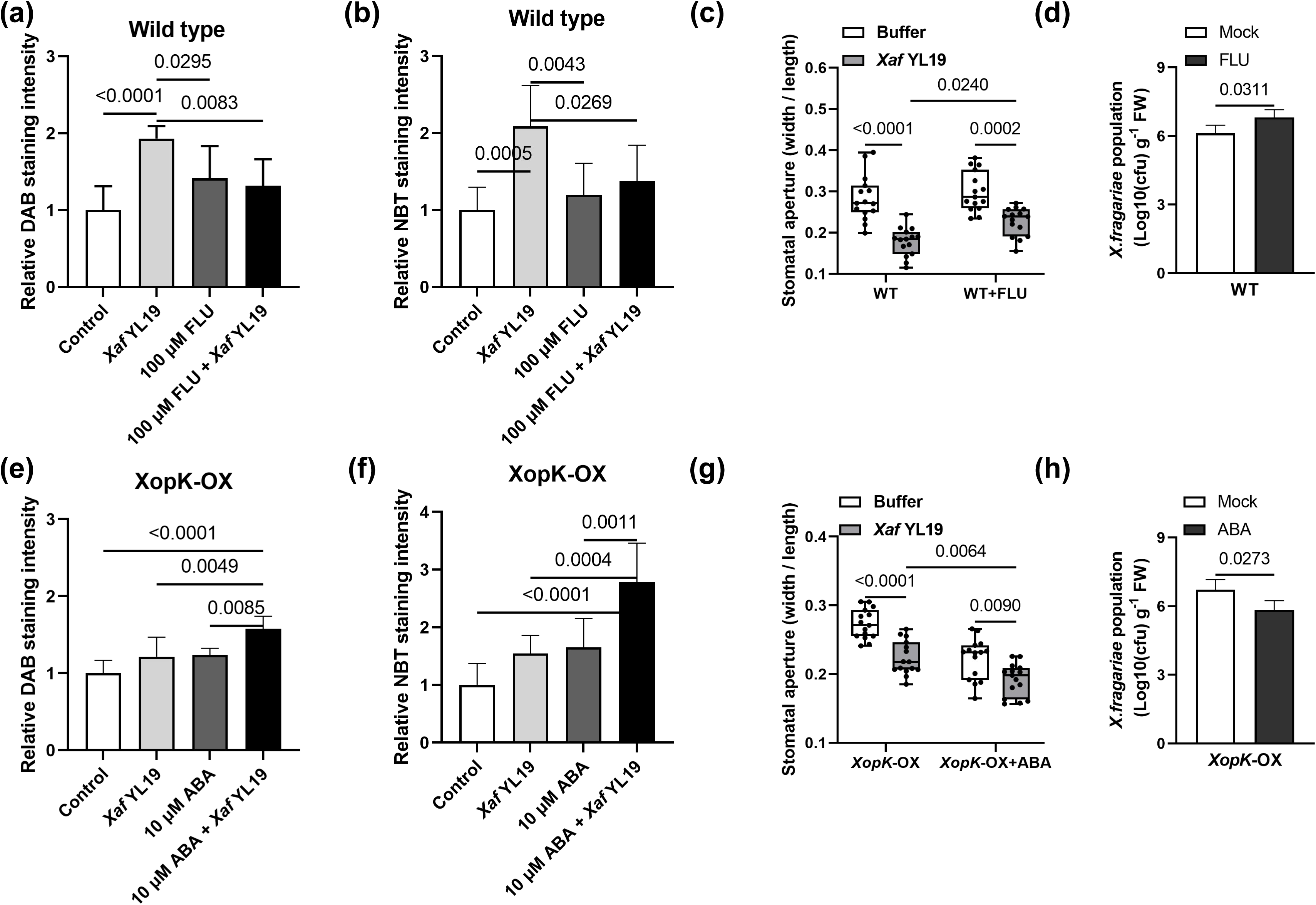
Effect of endogenous ABA on *X.fragariae*-induced reactive oxygen species (ROS) production and stomatal closure in strawberry. Fluridone-pretreating inhibited *Xaf*-induced ROS production (a), and stomatal closure (b), and bacterial population (c) in wild type of strawberry. ABA-pretreated increases XopK-OX strawberry ROS production (d), stomatal closure (e) and bacterial population (f). data are mean ± standard error, six leaves used for ROS measure in each treatment, The values on the top of each column indicate a significant difference by Two-way ANOVA (Dunnett’s multiple comparisons test).

## 3 Discussion

Type III effectors are important virulent factors in most pathogens, while there was limited knowledge of *Xaf* effectors and their virulent mechanism. In this study, we first identified a virulent effector XopK from thirty-three putative T3Es of *Xaf* YL19. XopK was initially identified as a putative T3E in *Xoo* due to the presence of a putative plant-inducible promoter box (PIP-box) (Furutani et al., 2006), later was observed to be secreted into host cells using a calmodulin-dependent adenylate cyclase (Cya) reporter system (Furutani et al., 2009), as we observed using phiLOV2.1 fusion fluorescence protein system (Fig. 1e). Then, the homolog XopK*_Xoo_* in *Xoo* PXO99A was proofed harbors E3 ubiquitin-ligase activity, and inhibits pathogen-associated molecular pattern-triggered immunity (Qin et al., 2018). We found that XopK completely suppresses plant cell death (Fig. S3). However, the homolog of XopK virulence depends on host-pathogen interaction. XopK*_Xoo_*-deletion strain of *Xoo* PXO99A affected in pathogenicity of the rice cultivar Nipponbare (Qin et al., 2018), but not in rice IR24 plants (Song & Yang, 2010). The XopK*_Xpm_* mutant strain of *Xanthomonas phaseoli* pv. *Manihotis* (*Xpm*) enhanced cassava disease symptoms at the inoculated site but limited the spreading through the vasculature (Mutka et al., 2016). The XopK*_Xcc_*-knockout strain of *Xcc* 8004 was not affected in pathogenicity on *Arabidopsis*, but ectopic expression of XopK resulted in *Arabidopsis* seedling growth inhibition (Noe Arroyo Velez, 2022). Considering more efforts needed on the gene-editing of *Xaf* (Vandroemme et al., 2013), as an alternative strategy, we generated the ectopic overexpression of XopK in strawberry, like *XopAP*-, *XopC2-* overexpressed in rice plants (Liu et al., 2022; Wang et al., 2021), evidenced that *XopK*-expressing lines exhibited significantly (*P*<0.05) higher bacterial replication at 4 dpi but not in a more later stage (8 dpi), previous work did not show the phenomenon in XopK*_Xoo_* transgenic Arabidopsis for *P. syringae* DC3000 *hrc*C-inoculation may because of different inoculation methods or statistic period (Qin et al., 2018). However, a more severe dry cavity rot in the crown at 45 dpi was observed (Fig. 3), which may account for the systematic moving of the early colonized bacteria from leaves to crowns through vascular tissues as described by Wang et al. (2023). The different infection outcomes with pressure or syringe-inoculation (Fig. S7), combined with stomatal aperture analysis during infection (Fig. 4) push us to focus on the function of XopK in stomatal immunity, which is one of the first lines of plant innate immune responses to most pathogenic bacteria (Zhang et al., 2020).

Functional ABA signaling is essential for stomatal closure during stomatal immunity (Melotto et al., 2006, 2017), and plants tend to keep higher basal ABA levels and ABA signaling in guard cells for maintaining steady-state stomatal conductance (Gonzalez-Guzman et al., 2012; Lahr & Raschke, 1988; Waadt et al., 2015). Hence, phytopathogen effectors often function ABA as a virulent target. They create a water-soaking microenvironment in the apoplast by inducing a strong ABA signature for stomatal closure (Hu et al., 2022; Roussin-Léveillée et al., 2022), or inhibit the biosynthesis of ABA in guard cells to open stomata that benefit initial infection (Liu et al., 2022). Here, we also found a significant decrease in basal ABA concentration and signaling in *XopK*-expressing strawberry plants (Fig.3). However, we do not dig into how XopK affects ABA accumulation and signaling in this study, it may directly suppress the expression of *FveNCED5* (Fig. 4b), which is the rate-limiting enzyme for ABA biosynthesis, seems like AvrPtoB, which promotes ABA accumulation in *Arabidopsis* by increasing the transcript level of *9-cis-epoxy carotenoid dioxygenase 3* (*NCED3*) in ABA biosynthesis (de Torres-Zabala et al., 2007), but latter proved its effect on the synthesis of ABA is most likely indirect (Cheng et al., 2011; Shan et al., 2008). The *Xoc* effector AvrRxo1, which reduces vitamin B6 (VB6) levels, and consequently inhibits the biosynthesis of ABA, provides an indirect case (Liu et al., 2022).

Like the ABA-deficient *aba2-2* mutant in Arabidopsis (Hossain et al., 2011), the application of 10 μM ABA significantly induced stomatal closure in *XopK*-OX strawberry, albeit at a delayed effect (Fig. 5a, b). MeJA did not induce stomatal closure in *aba2-2* mutants, a robust suppression of MeJA-induced stomatal closure in *XopK*-OX strawberry was also observed (Fig. 5c, d). However, that does not associate with the JA accumulation and signaling, as we evidenced that a wild type levels of JA and *FveJAZ1* expression, and even down-regulation of *FveLOX14* in *XopK*-OX strawberry lines (Fig. S7). Stomatal defense is often compromised when JA signaling is activated in the context of pathogen infections, as shown by two effectors XopC2 from *Xoc* RS105 (Li et al., 2014; Wang et al., 2021) and XopS from *Xanthomonas campestris* pv*. vesicatoria* (*Xcv*) (Raffeiner et al., 2022), but not in the XopK case. That suggests a more complex mechanism for the suppression of MeJA-induced stomatal closure by XopK. Previous studies showed signal crosstalk between MeJA and ABA in guard cells cannot be ignored (Munemasa et al., 2011; Zhu et al., 2012), MeJA-induced stomatal closure defective may be caused by the low endogenous ABA accumulation in *XopK*-OX strawberry (Fig. 4a), as an endogenous ABA threshold is needed for MeJA-induced stomatal closure (Hossain et al., 2011), also supported by the fact that stomatal movement of ABA-insensitive mutants is insensitive to MeJA (Munemasa et al., 2011). Whatever, the XopK transgenic strawberry may provide a good genetic material for cross-talk between ABA and MeJA in strawberries in future studies.

ROS signaling is often a convergent point of a second messenger for various pathways containing ABA- and MeJA-induced stomatal closing (Singh et al., 2017). We also found ABA and MeJA could induce ROS production in wild type by DAB and NBT staining (Fig. S10), while ABA- but not MeJA-mediated ROS production was defective in *XopK*-OX strawberry, like the ABA-deficient *aba3-1* mutant (Van Gijsegem et al., 2017). XopK may act ABA signaling upstream of the branch point of ABA signaling and MeJA signaling, and affect the downstream of ROS production in MeJA signaling. In addition, *XopK*-OX strawberries exhibit compromised bacterium-induced ROS burst and stomatal closure (Fig. 6). That is possibly attributed to a lack of ABA-induced ROS production. Similarly, *Pst* DC3000-triggered stomatal closing is affected in the *notabilis* tomato mutant, which harbors a null mutation in the ABA synthesis gene *NCED1* (Du et al., 2014). The above analysis indicated impairment of ABA-induced ROS production by XopK may be responsible for *Xaf* infection initially.

Based on our results, we presented a simple model of the *Xaf* YL19 infection under the favor of effector XopK in Figure 7. This model points out that during pathogen infection, the pathogen was detected by pattern-recognition receptors (PRR) complex localized in the plant plasma membrane, and function ABA and JA to trigger ROS production for fast stomatal closure and restrict bacterial invasion (Fig. 7, left model). However, the apoplast-colonized *Xaf* will secrete effectors XopK into host cells, then compromise ROS production via directly or indirectly inhibiting ABA biosynthesis and signal transduction or suppressing downstream of ROS production in JA signaling pathway, both of them make stomatal reopening and allow the pathogen to enter successfully (Fig. 7, right model). To the best of our knowledge, this study provides a new insight into the *Xaf* effector that manipulates hormone signaling to facilitate infection in strawberries.

**Figure 7.**
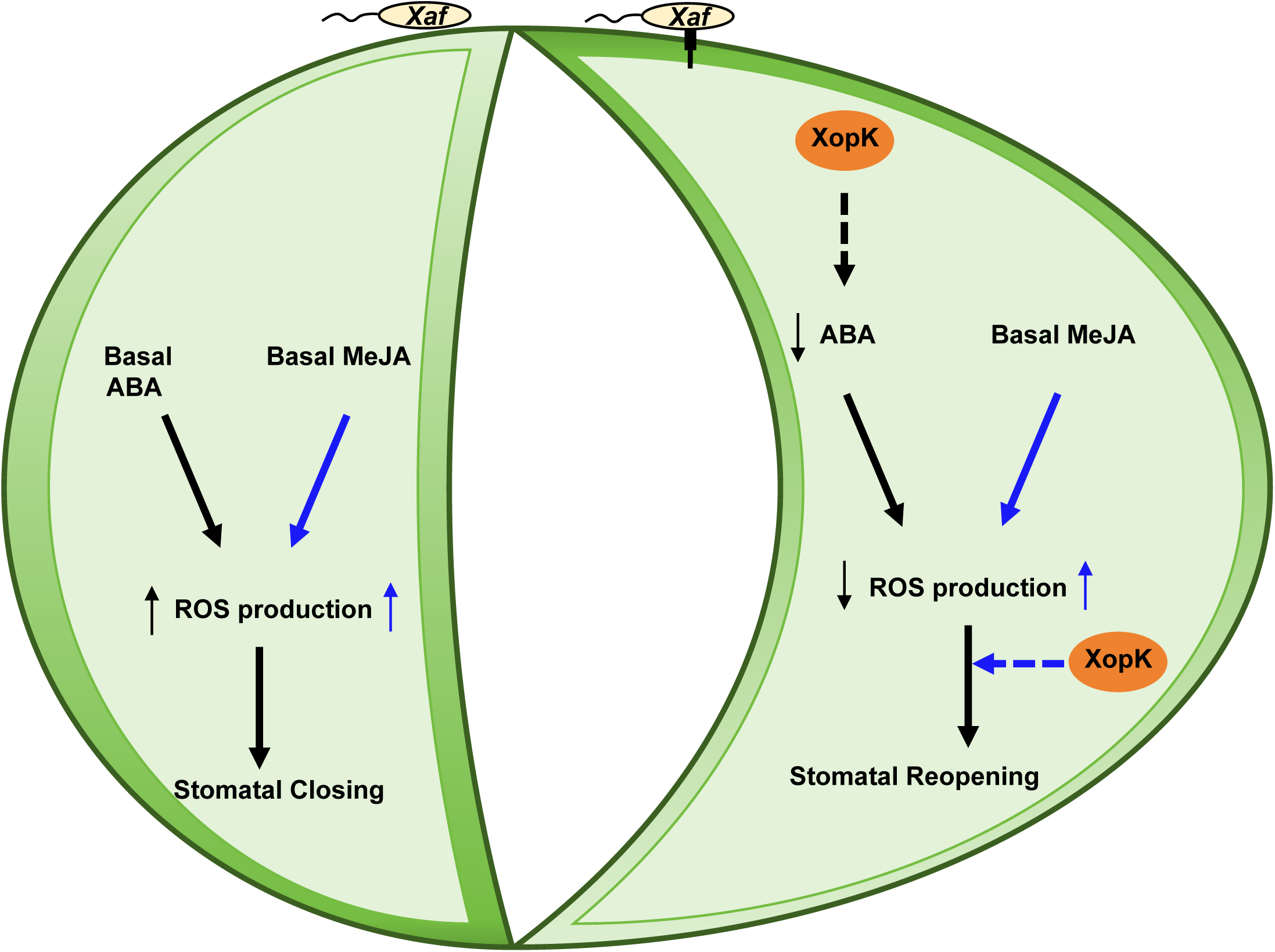
Proposed model for the function of XopK in stomatal movement of strawberry during pathogen infection. During *Xaf* infection, pathogen-associated molecule patterns (PAMPs) were detected by pattern-recognition receptors (PRR) complex localized in plant plasma membrane, and function ABA and JA to trigger ROS production for fast stomatal closure and restricting bacterial invasion (Left model). However, the apoplast-colonized *Xaf* will secreted effectors XopK into host cells, then may direct or indirect inhibit ABA biosynthesis and signal to compromise ROS production, and suppresses downstream of ROS production in JA signaling, both of them make stomatal reopening and allowing the pathogen to enter successfully (Right model). Black and blue lines indicate ABA and MeJA effects, respectively, and solid and dotted lines indicate direct and indirect action. respectively.

## 4 Materials and methods

### 4.1 Biological materials and growth conditions

*Xanthomonas fragariae* strain YL19-GFP is grown in liquid nutrient broth (NB) or NB with agar at 25 °C (Wang et al., 2023). *Escherichia coli* DH5α was cultivated at 37°C in either liquid or solid Luria-Bertani medium. The transformants *Agrobacterium tumefaciens* GV3101 were cultivated at 28°C. For this assay, antibiotics were used at doses of 50 μg/mL for kanamycin, 100 μg/mL for ampicillin, 50 μg/mL for rifampicin, and 50 μg/mL gentamycin.

*Fragaria vesca* ‘Hawaii 4’, and *Nicotiana benthamiana* plants were grown in a controlled environment with exacting conditions: 25°C temperature, 16/8 h light/dark cycle, and 75 % relative humidity (RH).

### 4.2 Plasmid construction and plant transformation

For expression of phiLOV2.1-tagged XopK protein in *X. fragariae* YL19, The XopK gene from the genomic DNA of *Xaf* YL19 was amplified by polymerase chain reaction (PCR). The synthesized DNA fragments of phiLOV2.1 (Tsingke, China) were tagged to the C-terminal of XopK and subsequently ligated into pBBR5 containing the *XopK* promoter (960 nucleotides upstream from the translational start codon) using *EcoR*I and *BamH*I producing pBBR5[*XopK:phiLOV2.1*], as described by (Roushan et al., 2018). The phiLOV2.1 was only cloned into pBBR5 under the control of the XopK promoter, creating pBBR5[*phiLOV2.1*] as the negative control. Both pBBR5[*XopK:phiLOV2.1*] and pBBR5[*phiLOV2.1*] were electro-transfers into the competence of *Xaf* YL19 and cultured in NB medium containing gentamycin at 25 °C.

For examining plant cell death suppression, the code sequence of T3E genes was cloned into the potato virus X-based vector pGR106, then electro-transfer into GV3101 (containing helper plasmid pJIC Sa_Rep). The Agrobacterium containing vector pGR106[*Avr1b*] and pGR106[*BAX*], which were gifted by Professor C.L Tang, were used for positive control in suppress and induced cell death assay, respectively. For the assay of subcellular location and transgenic plant, the DNA fragments of *XopK* (cloned from *Xaf* YL19 DNA), and *AtPIP2A* (cloned from *Arabidopsis col*-0 cDNA) were ligated into the pCAMBIA2300-GFP with the regulation of the cauliflower mosaic virus (CaMV) 35S promoter. Both of them were transformed into Agrobacterium GV3101. The corresponding Agrobacterium was used for the generation of transgenic strawberry based on the method described by Ma et al. (2023).

### 4.3 Agrobacterium-mediated transient gene expression in planta

The infiltration solution (10 mM MES, 10 mM MgCl_2_, and 150 μM acetosyringone, pH 5.6) was used to dilute the cultured Agrobacterium to an optical denseness at 600 nm (OD_600_) of 0.2, which was then introduced into *N. benthamiana* with a 1 mL syringe. For co-infiltration experiments, different recombination vectors were carried by *A. tumefaciens* in a 1:1 ratio with infiltration buffer, resulting in an end optical density (OD_600_) of 0.2. The leaves were gathered for further microscopy 2 days after infiltration or used for observation and analysis of plant cell death 5 days after infiltration.

### 4.4 Virulence assay

The virulence of the *X.fragariae* YL19-GFP strain in strawberry plants was determined based on previous studies (Liang et al., 2023; Wang et al., 2023; Wei et al., 2024). The YL19-GFP was cultured in liquid NB medium to OD_600_=1.0 (10^8^ cfu/ml), then resuspended in 10 mM MgCl_2_ buffer with 0.01% Silwet L-77 for subsequent inoculation assay. Following inoculation, the plants were kept in a growth chamber. Each replicate assay with twelve strawberry plants and the disease index (DI, %) in leaf and incidence of dry cavity rot in the crown (IDCR) was calculated at 8 dpi and 45 dpi, respectively, with the following formulas (Wei et al., 2024): DI (%) = (Sum of individual ratings / Number of plants examined × Maximum disease scale) ×100; IDCR (%) = (Number of plants with dry cavity rot / Number of inoculated plants) ×100. For the leaf disease index, two fully expanded leaves of each plant were taken photo and the leaf lesion area was calculated using ImageJ software (http://imagej.nih.gov/ij/).

The biomass of YL19-GFP was measured by Real-time PCR as described by Wang et al. (2023). Leaf samples were collected at 0, 4, and 8 dpi, then gently washed with sterile water and frozen liquid nitrogen for DNA extraction and RT-qPCR analysis.

### 4.5 Stomatal aperture

Stomatal aperture assay was carried out as described by Shang et al. (2016). Three-month-old strawberry leaves were incubated for three hours under intense light in stomata opening buffer (30 mmol/L KCl, 10 mmol/L MES-KOH, and 10 mmol/L CaCl_2_, pH 6.05) to completely open stomata. Subsequently, the leaves of the strawberry plants were subjected to 10 µM ABA, and 75 µM MeJA. Stomata images were captured using a confocal laser scanning microscope (Leica TCS SP8 SR, Germany) with exciting by 405 nm laser for auto-fluorescence from the inner cell wall of guard cells. For each treatment, a minimum of 20 stomata were chosen at random. ImageJ software (http://imagej.nih.gov/ij/) was used to measure the aperture size.

### 4.6 Water loss assay

Detached leaves of strawberry plants at the same leaf old were kept in a growth chamber, and their initial weights were weighed individually. Next, weigh for the water loss every ten minutes over a 40-minute course.

### 4.7 Phytohormone measurements

Strawberry leaves (50 mg) were ground into powder in liquid nitrogen using a high-throughput tissue grinder (Scientz, China). The mixture was shaken for 10 minutes at 30 HZ using a tissue grinder after adding 1 mL of ethyl acetate. The supernatant was transferred into a new 2 mL centrifuge tube after centrifuging at 12,000 rpm for 10 minutes. The sample was spined in a centrifuge at 13,000 rpm for 15 minutes after evaporating the ethyl acetate with a nitrogen blower until it was dry.

Then, add 200 μL of 50% methanol to the sample that has evaporated. 100 μL of the supernatant was filtered through a 0.22-μm filter for Liquid Chromatography-Tandem Mass Spectrometry (MS/MS) analysis.

### 4.8 RNA isolation and quantitative RT-PCR

Three-month-old strawberry plants were sprayed with 1×10^8^ cfu/ml *Xaf* YL19 suspension, and bacterial cells from strawberry leaves were gathered at 0, 6, 24, 54, and 96 h post-inoculation, and extracted RNA using the Bacterial RNA Extraction Kit based on the manufacturer’s recommendations (Vazyme Biotech, China). Plant RNA was extracted by E.Z.N.A. Plant RNA Kit (Omega, BIO-TEK, USA). First-strand complementary DNA (cDNA) was synthesized with TransScript^®^ One-Step gDNA Removal and cDNA Synthesis SuperMix (TransGen, Beijing, China). An Applied Biosystems^TM^ StepOnePlus^TM^ Real-Time PCR System (ThermoFisher, USA). The gene-specific primers are listed in Table S2. For the normalization of *Xaf* YL19 and strawberry genes, the reference genes *pykA* (NCBI GenBank ID: CP071955.1), and *FveGAPDH* (NCBI GenBank ID: XM_004287072.2) were used, respectively. The TBtools software was used to estimate the fold change in gene expression (C. Chen et al., 2020).

### 4.9 Stomatal conductance measurement

Stomatal conductance in leaves of three-month-old strawberry plants using a photosynthesis system (LI-6400XT, USA) with a leaf chamber (2×3 cm) equipped with a red-blue LED light source. After exposing the leaves to light in the growth chamber for 2 hours, they were inoculated with *Xaf* YL19. Stomatal conductance was then evaluated at 2 hours post-infection. Measurement parameters were adjusted to 400 μmol mol^-1^ CO_2_ and 200 μmol m^-2^ s^-1^ photosynthetic photon flux density.

### 4.10 Reactive oxygen species (ROS) staining

ROS level was detected by staining with 3,3‵-diaminobenzidine (DAB) and nitroblue tetrazolium (NBT) as described by Jambunathan (2010). For ABA- and MeJA-induced ROS assay, detached strawberry leaves merged into a stomatal opening buffer for 2 h, then incubated with 10 µM ABA or 75 µM MeJA for 30 min, the vacuum infiltrated with DAB solution and NTB solution overnight. The leaves were removed chlorophyll by 95% ethanol. For the *Xaf*-induced ROS assay, 100 µM fluridone or 10 µM ABA was pretreated for 3h in wild type and *XopK*-OX strawberry, respectively, which was followed by spraying inoculation with *Xaf* YL19 suspensions. ROS in whole leaves were visualized as a brown color by DAB staining or blue color by NBT staining, then quantified using ImageJ software (http://imagej.nih.gov/ij/).

### 4.11 Statistical analysis

Data significant differences were evaluated by the IBM SPSS software (v.26.0; IBM, USA). Figures were drawn using the GraphPad Prism 9 software (San Diego, USA).

## Credit authorship contribution statement

Xiao-Lin Cai: Data curation, Methodology, Formal analysis, Writing – original draft, review & editing. Wen-Yao Zhang: Methodology, Formal analysis–review & editing; Hai-Yan Yu: Investigation, Methodology, Formal analysis–review & editing. Ying-Qiang Wen: Conceptualization, Project administration, Funding acquisition. Jia-Yue Feng: Conceptualization, Project administration, Funding acquisition.

## Acknowledgments

The authors would like to thank the anonymous reviewers for their comments on the manuscript. This work was sponsored by the Key Technology Tackling Project of Agricultural Key Industry Chain in Xi’an (No. 22NYGG0006), a major project of Agricultural Cooperative Innovation and Extension Alliance in Shaanxi Province (No. LMZD202102), the Key Research and Development Program of Shaanxi province (No. 2023-YBNY-083), and the Scientific and technological innovation and achievement transformation project of experimental demonstration station (No. TGZX2021-20).

